# How researchers experience the impact of consortia and ERC funding schemes on their science

**DOI:** 10.1101/2022.07.30.501782

**Authors:** Stephanie Meirmans, Herman J. Paul

## Abstract

Policy makers push for consortia science geared towards addressing important issues. Such consortia are expected to target societal problems, be international, to engage in trans- or interdisciplinary research, to involve stakeholders and have specific plans for implementation. For example, Horizon Europe focuses on five missions that are being targeted by such type of consortia. This, however, does not seem to be the type of funding that active researchers appreciate the most: a recent letter signed by over 24.000 researchers clearly shows their preference for ERC grants. What are the underlying reasons for this difference? Here, we share insights on how natural science and medical researchers experience the impact of these funding schemes using interviews. Our findings highlight that the two different types of funding schemes have a different performative effect on research, and that ERC-type funding aligns most with how scientists think research should best be conducted.

## Introduction

Funding agencies spend considerable sums on fostering collaborative science (Wager 2018). In consortia funding, scientists typically target a specific societal problem or challenge in bigger interdisciplinary and often also international groups, working closely together with stakeholders and citizens. An international frontrunner for such consortia funding efforts is for example the EU in its top-down pillars of the Horizon 2020 framework (De Rijcke and Wilsdon 2019), but also national funders increasingly fund these types of consortia (for example the Dutch and Swiss national funders).

Against this trend in science policy, researchers themselves seem to value other, perhaps more traditional, types of funding schemes more highly. For example, in a recent ‘letter of the friends of the ERC’, over 24.000 researchers had signed a plea for not reducing funding towards the ERC (see Friends of the ERC, letter, https://friendsoftheerc.w.uib.no/the-letter/). What do such pleas mean for the current predominant trend away from more traditional types of funding towards even more problem-driven consortia types of funding science? And why exactly do scientists see that trend as a problem?

Notably, also a recent science studies paper (Falkenberg et al. 2022) urged national funders to stop modelling national schemes after the ERC. More specifically, these authors focus on the ERC as promoting a normative regime of innovation, of breakthrough science. Against this, and leaning on empirical evidence of several case studies, the authors argue that such innovation funding schemes only work if they are in a healthy balance with funding schemes that foster more incremental types of science. They called this the ‘breakthrough paradox’: too much funding towards breakthrough science will impede breakthrough science in the end. Also Scholten et al. (2021) have argued to reduce funding towards excellence schemes because they foster too much competition. These authors suggest rather providing funding to other types of funding schemes.

In this study, we report upon findings from a recent group interview study that provides epistemic reasons for strengthening ERC-type funding instead of consortia-type funding. We find that researchers prefer ERC-type funding not per se due to the innovation or excellence component, but because several aspects of the funding specifics align mostly faithfully with how they experience science should effectively be done, also in terms of impact: (1) in flexible and small-scale types of teams that focus on close collaborations and where team members can be added as seems most valuable to the science conducted rather than in loose networks that suffer from all kinds of frictions; (2) curiosity-driven rather than focusing on generating short-term impact that is experienced as highly unrealistic; (3) being autonomous and flexible in terms of choice of topics and methods rather than heeding to pre-structuring via funding calls that is sensed as complicating matters.

## Details of funding schemes

### ERC

ERC grants are essentially excellence schemes, meaning that they should provide the most talented scientists with money to pursue their ideas. These grants seem to have partly be modeled after US NSF research funding as well as after national excellence scheme precursors in the Netherlands, the so-called ‘Veni, Vidi, Vici’ schemes. ERC grants have a similar three-step funding scheme as the ‘Cesarian’ excellence schemes, going from smaller to bigger grants with a scientists’ seniority. ERC grants are also explicitly coupled with a notion of breakthrough science, which often goes under the header of ‘high-risk high-reward’ funding. Basically, the idea is to not only provide excellent scientists with the money to do their research, but these scientists are also supposed to follow up on daring ideas, pushing the boundaries of science, innovate. Philosophically, the ERC seems to be built on a Kuhnian idea of revolutionary (versus normal) science (see also Falkenberg et al. 2022). An ERC grant should provide scientists with the possibility to make that ‘big leap’ away from normal science. Typically, the scientist gathers a team to do the proposed research.

### Consortia science

The development towards consortia science is often justified with the philosophical argument that current scientific and societal problems are sufficiently complex to require multi-dimensional expertise, and that they can therefore only be effectively addressed by large teams of scientists with different disciplinary backgrounds and the involvement of potential stakeholders (Wickson et al. 2006; Falk-Krzesinski et al. 2011; Milojević 2014; National Research Council 2015; Cundill et al. 2019). This is a theoretically attractive idea because any scientific or societal problem can then be addressed from multiple perspectives, in the hope to thus overcome any potential biases stemming from a specific discipline or focus that are thought to hamper progress, and to include all relevant factors and aspects. Such an approach could also increase chances of realistically developing and implementing any needed changes and interventions. And indeed, science scholars describe that inter - and transdisciplinary approaches are characterizing contemporary science (as opposed to post-war II fundamental science; see e.g. Gibbons et al. 1994).

Many science scholars actively promote the funding of such types of bigger collaborations, across disciplines and with stakeholders. Amongst science scholars, this way of doing science goes under a variety of names today, such as ‘Mode 2’ science (Gibbons et al. 1994), transdisciplinary science (Wickson et al. 2006), post-normal science (Funtowicz and Ravetz 1990), post-academic science (Ziman 2000), knowledge co-production (Bremer and Meisch 2017), knowledge co-creation (Regeer and Bunders 2009), and (if it involves industries and universities) triple helix relations (Etzkowitz and Leydesdorff 2000). Finally, recent approaches often focus on RRI (Responsible research and innovation) concepts, which specifically aim to more flexibly integrate the science-society divide (Owen et al. 2012).

Perhaps not unimportantly, funding consortia science often goes well with politics, science policy and citizen engagement. By covering many dimensions, funding agencies can meet prevailing high standards of accountability via not leaving out any important factors or (political and social) dimensions. Finally, such type of science funding often explicitly focuses on immediate public needs. For example, Horizon Europe missions include fighting cancer and climate change, work towards cleaner oceans, waters, coasts and soils, as well as promote greener energy (Wallace 2020).

## Science studies on ERC and consortia science

Overall, ERC and consortia science seem to be built up on two different types of epistemologies: break-through science (ERC) on the one hand, and co-creation science (consortia) on the other hand. The ERC is internationally seen as a big success story (European Research Council 2019). A recent science study has shown that such excellence grants can indeed provide researchers with the resources to do significant work and give them epistemic and organizational autonomy (Scholten et al. 2021), though this is even more so the case for prize funding (Franssen et al. 2018). On the other hand, Scholten and colleagues have also shown that even if researchers have an excellence grant, there is a constant need to compete for future grants, a state they call ‘strategic anticipation’. Due to this competition, coupled with the fact that only few groups can benefit from excellence funding, Scholten and colleagues argue that it might help ‘to decrease the budget for excellence funding arrangements, allocating the rest of the funding to other funding programs or as block funding’. Also another science study has recently pushed the idea that excellence funding might not work well in practice, but due to another reason. Falkenberg (2021) found that funding schemes like the ERC with their focus on innovation can be in tension with at least some scientific practices towards how innovation works because also breakthrough science always needs normal science as a base. Falkenberg et al (2022) therefore argue that it would impede scientific breakthroughs in the end if all funding would be geared away from normal science towards breakthrough science in an ERC-style. Falkenberg and colleagues urge for a better balance between innovative and incremental science in the funding ecosystem.

Many science scholars are convinced that knowledge co-creation, on the other hand, does work well. Some have argued that it is socially responsible to push through a mode-2 type of science – even against the preferences of the researchers themselves (see e.g. Chubb and Reed 2017). What is needed if this creates friction is to educate scientists to value broader contexts and become more reflexive (e.g. Åm 2019), and/or to better align the incentives of other stakeholders that can play a role in efforts towards more socially responsible science, such as universities (Sigl et al. 2020).

Science studies researchers investigating actual efforts of transdisciplinary research in practice, however, tell a somewhat different story. Across different domains of research practice, they all mark how difficult it is to do such type of research well. For example, while Ribeiro et al. (2019) remain convinced of the benefits, they also point to a host of challenges and problems inherent to transdisciplinarity in One Health research. They find high administrative and managerial burdens and major organizational and integrational challenges linked to the diversity of perspectives and power relationships involved in larger teams (see also Pohl and Hirsch Hadorn 2008). Likewise, researchers working on sustainability issues find that knowledge co-production between researchers and non-academic stakeholders can be very complex. Recent research in this area suggests that knowledge co-production does not always lead to positive effects and that far more research is needed to determine under what conditions knowledge co-production is effective and desirable – and when it would be better to abstain from it (Lemos et al. 2018; Wyborn et al. 2019).

The National Research Council report (2015) urged that more research would be needed to understand how alternative funding strategies may affect team science effectiveness. Funding aimed to bridge the science-society divide, such as ELSA and RRI funding, has recently been shown to also have problems and trade-offs once being put into research practice (e.g. van Hove and Wickson 2017; Carrier and Gartzlaff 2019). It remains to be seen how the specifics of consortia science and ERC funding affect (team) science in practice.

## Empirical study: Methods and details

Findings for this paper were extracted from interview sessions which we conducted in 2017/2018 in the Netherlands and in Switzerland. In this research, we explored how active scientists experience and perceive the impact of competitive research funding on their science. This paper focuses on those statements and comments that can help to gain an understanding of the performative effects of both ERC and EU consortia funding schemes on scientific practice.

### Session participants and details

For our research, we had conducted six group session interviews in two countries. The groups consisted of three to seven researchers, grouped by scientific domains (natural sciences, medical sciences, or humanities) and career status (junior = holding a temporary job position, or senior = holding a permanent position). This made a total of twelve group session interviews with in total 57 persons. Interviewees were recruited via personal networks as well as via Dutch and Swiss university websites and the website of the Royal Netherlands Academy of Arts and Sciences. This recruitment strategy resulted in a significant number of very experienced researchers, also with large-scale (consortia) funding, in particular amongst the permanent staff. Each session took around 3.5 hours. We taped the oral discussions and subsequently had them transcribed by a professional transcription bureau.

We also used ‘Meetingsphere’, a tool designed to allow anonymized digital interaction between the group members (https://www.meetingsphere.com). The interviews revolved around the themes of how competitive research funding affects science and how funding could be improved to foster good science. Group members were first allowed to type their comments into the digital system. After saturation of commenting (typically after 10-15 minutes), we opened the system up for digital commenting, followed by extensive oral discussion. One of the groups ended up with oral discussion only (due to the delayed arrival of one participant).

### Analysis

We used a thematic analysis to analyze the transcribed interviews and Meetingsphere reports. We here present exclusively the results concerning the interviewees’ perceptions of EU-consortia and ERC funding^1^. Where it helps to understand the issues at hand, we also report on experiences with other national forms of funding (e.g. Dutch excellence funding and other forms of national consortia funding). We also made the decision to exclude the humanities from this paper as experiences in this field diverge too much from experiences in the medical and natural sciences and will therefore form a separate future paper. Regarding experiences in the natural and medical sciences, we basically found that there are three different themes connected to the ERC and consortia funding schemes that we present in detail here: type of collaborations, purpose of funding and organization of funding.

## Results

### Type of collaborations: one PI and a close team versus a loose consortium network

Our interviewees emphasized that one of the aspects in which ERC and EU consortia funding differ is in the type of collaboration, which is not surprising given that this is indeed one of the core differences between the two funding schemes.

The ERC is granting ‘hard-core personal subsidies’ (med sen CH), meaning this is a grant awarded to a single person. This PI then can (and typically does) gather a team around him/her who can co-work closely in the project. One Swiss senior medical researcher thought that ERC grants are phantastic, they are ‘kind of an award’, even though ‘the acceptance rate is going too low’. Across all the focus groups, we have not heard one researcher saying that the collaborative structure within an ERC grant structure was not working out as intended. This might not be so surprising as such teams are flexible in size, put together by the PI and eventually work closely together within one institution. Other science studies have found that working closely together makes for the most easily-achieved successful types of collaborations, while looser types of networks need well-coordinated organization, including physical getting-together, to become successful (Hesjedal 2022). Also the social dynamics are found to be highly important for success within a collaboration (Dusdal and Powell 2021; Hesjedal 2022).

In contrast to a personal grant such as an ERC, a consortium typically consists of a loose and large network of researchers across universities. Many interviewees across natural and medical science groups acknowledged that consortia are ‘a good incentive for collaborative research’ (med sen CH). However, the general tendency of our natural and medical senior researcher interviewees who had experience with such funding was that they are not too happy with the resulting type of collaboration. One big issue was that the collaborations ‘often do not work very well-communications issues-between disciplines and need better support and guidance’ (med sen CH). And indeed, that such communication problems frequently occur in international big collaborations has been reported elsewhere (Dusdal and Powell 2021).

In general, researchers thought the bigger the consortia the worse they work in practice: ‘my subjective personal experience is that the larger the consortia are, the smaller is the input/benefit ratio’ (med sen CH). And a senior Dutch medical scientist said ‘I can’t say that I found the research better than the sum of its parts. In fact, it was worse…’. Similar to EU consortia, also for Swiss NCCR’s (National Centres of Competence in Science) which are national bigger types of consortia across 10-12 collaborators, Swiss scientists experienced problems if too many people became involved; it resulted in that ‘you try to avoid meetings, because somebody constantly leaves the lunch meeting’ (med sen CH).

In a similar way than the medical researchers, Swiss natural scientists emphasized that bigger consortia might not always serve their purpose. For example, when one Swiss natural scientist emphasised the positive effects of funding because it ‘may push scientists to interact [via collaborations] and accelerate discovery’, another scientist immediately went against this: ‘True if it works that way. However, for large scale networks this may also lead to a lot of formal interaction without actual benefits’. Indeed, they experienced that EU Horizon projects were ‘more on paper than real’. Medical scientists experienced this in a similar way: ‘It was not really a true collaboration. It was just opportunistic that people found each other, because they knew they could get easier money that way.’ And that if you ‘write it down in a nice way, then it looks fantastic, but it’s an empty bubble, really.’ (both quotes med sen NL)

In addition, it was also not clear how success in bigger consortia structures should be assessed. For example in connection with NCCR’s, one natural junior Swiss scientist said: ‘what do you harness as a success? That two of these centres somehow connect, or is it only a success if all of them connect and form a single structure, or is it just a success if you get two or three more links between them?’ The same reasoning holds for the bigger EU collaborative projects, where positive effects of networking were experienced but overall there were doubts that making such networks is worth the money: ‘It’s also the same for the European ones. I guess, I mean, the networks will perhaps not re-establish long-term structures, but they will perhaps still establish many small links. This may or may not-it’s certainly good, but I’m not sure if it’s…worth the money.’ (nat jun CH)

Another Swiss natural science junior researcher had personal experience with bigger consortia in the UK and was not impressed how these big consortia worked out in practice in that country. He/she called such consortia a ‘galactic waste of money’ and said that ‘a huge network just for the sake of making a huge network, I don’t see the point. It feels a little bit showering money down to academia just so that everybody has something to do’. In addition, ‘there have to be administrators. I don’t know, I don’t think it’s a good way…’. And again, other studies have pointed out that excessive administrative work can be a problem in such large-scale collaborations (Dusdal and Powell 2021). Our interviewee emphasized instead the need to have funding for smaller interdisciplinary collaborations, not ‘gigantic things’ but instead ‘to work with a colleague’.

Swiss medical seniors also pointed out that another problem with such big consortia can be what kind of contribution you would want to do in such a big structure. That even though ‘the scope can be very ambitious…that doesn’t mean that within the consortium, you are doing the most ambitious contribution.’ It seems that this may have to do with ownership of scientific ideas and insights or perhaps because other members of the consortium may think in different ways. Again, this has been reported from other international large collaborations as well, and Dusdal and Powell (2021) therefore urge to make clear authorship deals and/ or be flexible with who is allowed to publish what.

Another related issue that our interviewees reported upon is that many researchers want to become part of a consortium (due to the money involved), even though their inclusion might have a negative effect on the overall outcome: ‘the larger, [….] the higher the danger is that many people are pushing themselves into such a construct, just bending their expertise a little bit in order to fit in. And there’s a lot of friction and constraint and loss of resources into that part’ (med sen CH). Again, also Dusdal and Powell (2021) urge that members in international big collaborations should be picked out well to align scientific backgrounds and contributions, and to reduce frictions due to communication problems or different research or epistemic cultures. In comparison, other types of collaborations do work well, Swiss senior medical scientists emphasized, for example the smaller interdisciplinary Swiss Synergia projects. These projects are small consortia of three to four members. You ‘can really work together in a different field. It makes sense. Creates community. Creates interaction.’ It is likely that such smaller constructs are largely able to avoid the frictions and problems that have been identified for larger networks and can therefore harness successes more quickly and easily.

Several Dutch natural science seniors emphasized the problem that often high-achieving and visible researchers get invited to become a part of such consortia; these are, they say, always the same ‘usual suspects’. Such Matthew effects (alluding to that those researchers that have successes will easily gain more successes; Merton 1968) are well-known career effects in science and have already been described for gaining funding. Our interviewees here suggest that this effect also plays a role for who gets invited as collaborators. In fact, including such high-flyers would make strategic sense to increase success of getting the funding. Our Dutch senior natural science interviewees in any case got so annoyed by this effect that several of them have started to rather include researchers as collaborators that are fun to work with. Two scientists had independently put together proposals with such an idea in mind: ‘who would we actually want to work with?’ Rather than inviting the ‘usual suspects’, they invited ‘not the people who you see have the highest publication record, but just people that can take this challenge and think beyond, really thinking out of the box, are creative and team workers, and all these other skills, and who are nice people to have around because if you’re locked up in a room for a week you have to like them…’ (nat sen NL). Interestingly, recent science studies suggest that this may be a strategy that could pay off extremely well in practice: Dusdal and Powell (2021) recommend to not neglect the social factors that matter for successful collaborations, such as being friends (see also Hesjedal 2022).

### Purpose of funding: Curiosity-driven versus impact-driven science

ERC grants are curiosity-driven bottom-up grants, supporting fundamental science, while consortia funding schemes are geared towards generating societal impact. We did receive many positive and absolutely no negative statements regarding the purpose of funding fundamental science with ERC grants, while there were plenty of negative comments regarding impact-driven science, across the interviewed medical and natural science groups.

One Dutch medical senior scientist emphasized that team science as funded by the ERC does function well, basically because there is no other agenda than the science itself behind it: ‘because those are hard-core, personal subsidies with no commercial interests whatsoever…that’s why group science really flourishes there. There’s no financial or economic agenda hidden behind it. Whereas, for all others, there are these considerations.’ Also Swiss medical senior scientists emphasized that the ERC is very different from other EU programmes because the ERC is scientific research while many other programmes do have a secondary objective. For example, when regarding Horizon 2020: ‘this is research to which a lot of other things have been loaded on top and then this is, sort of, a very mixed bag, which I think is very different from the ERC, which is a research project.’ (med sen CH)

Two Dutch medical seniors also emphasized that this positive effect is partly also true for the Dutch excellence grants – at least once you have these types of grants they are pretty flexible, ‘you can just spend the money on what you want’. The underlying reason for why they say that you first need to get them and only then you are free to do as you wish may relate to the fact that also Dutch excellence grants are judged by their lengthy valorisation section during proposal peer review (de Jong 2015; Brenninkmeijer 2022).

Senior medical Dutch researchers all agreed that EU-Horizon proposals are driven too much by impact. One highly experienced researcher with many grants, including Horizon 2020 grants, even claimed that they are ‘hot air’; they promise impacts that cannot be realized, and ‘those proposals are empty proposals’. Dutch medical researchers differed whether the underlying science base needs to be good or not, though. But they did all agree that the time frame is completely off: ‘…they use these words like, “We will abolish dementia from the world” and things like that…I’m like, be realistic, within three or four years, you’re not going to do those things.’ This may be due to ‘the economic push, people want to see a return of investment in three or four years.’ However, this is highly unrealistic in medical realities even when considering medicine that is now considered highly successful: ‘the proper timeline for return of investment should be, at least, 15 years to 30 years. Not three years. It’s unrealistic. So, I don’t know why they require you to write that in the proposal.’ (med sen NL) Like their Dutch colleagues, medical senior scientists highlighted that the expected impact time horizon often is highly unrealistic (here: regarding Horizon 2020 consortia): ‘You know, they make programmes to find new a new drug against depression in five years. I mean they will just fail, there’s no question.’ (med sen CH)

One Swiss senior medical researcher thought that it would be important for funding agencies to understand how scientific breakthroughs work in practice: ‘I think every major scientific breakthrough has come out of probably some surprised discovery that was completely unplanned. From, probably, people who were looking for something else.’ (med sen CH) This means planning in the direction of impact, and for sure short-term impact, is probably not working, according to this scientist. What is more, scientific findings that did not seem important at first can lead to a breakthrough decades later. Other medical scientists alluded to the same problem: ‘If something is predictable, you cannot call it research.’ (med sen NL; with other colleagues agreeing to this).

The Swiss senior medical researcher added that it would therefore be ‘important to tell the public that they have to be able to tolerate a huge amount of…not successful experiments and labs.’ And that ‘they have to understand that really big discoveries were made not with plans to make them.’ The only thing that would help, according to this medical scientist, is to hire people with a certain personality, who are ‘curious and diligent and, actually, really follow up things’. So essentially ‘for the system to work…there has to be a certain basis of rewarding such, literally, playing around’. And one could historically argue that ‘every major discovery has come from such type of behaviour’. It is interesting to see here that this researcher essentially makes a move from a focus on impact to a focus on the scientist as a person, and that this researcher thinks even in terms of long-term impact it would be more valuable to fund persons (as in an ERC grant) rather than impact-driven consortia.

Medical senior researchers also talked about problems with stakeholder involvement required by Dutch funding agencies. In particular the currently often-made requirement of co-funding (meaning that stakeholders are supposed to also co-fund the research), ‘is extremely limiting’. It in practice inhibited this researcher to submit an important grant proposal because it could not be realized before the end of the submission deadline. This researcher complained that ‘I don’t know why they make such a strong requirement.’ Another researcher thought that the underlying reasons are ‘to show that those parties are really interested and willing to give money and, also, give time and effort to it I think. At least, in our field, it’s not so much industry but local care companies or whatever.’ (med sen NL) Interestingly, when asked what this researcher thought about involving care companies, the researcher said that ‘it only makes things more complex. And doesn’t necessarily say so much about how interested they [the local care companies] are…It can, more, be a hurdle than something else…’ Several other medical researchers in the group agreed with this. This researcher then also questioned whether the fact that such companies need to pay money would make it ‘more valuable for them.’ This is interesting in connection with co-creation ideas: our interviewees essentially experienced that involving stakeholders does in fact not help but rather complicates the research process.

Dutch medical senior researchers also added that the Dutch medical funder ZonMw does not merely expect co-creation but also expects a firm plan for implementation. However, this can be premature: ‘you also get that you don’t know, yet, if something works, but you already have that plan for implementation.’ And what is more, this all complicates the research process to a degree that the core research cannot be attended to in a manner that it would deserve, for example because budget needs to go towards implementation. It thus seems that at least for Dutch ZonMw grants, added aspects of co-creation and implementation in short-term research funding schemes may have a negative effect on the research as well as even on the eventual societal impact. Funder plans are experienced as over-ambitious, with several Dutch senior medical researchers sharing this experience.

Natural scientists had other worries than their medical colleagues: they mostly worried about a push away from basic research towards applied research via the need for funding. Indeed, only one researcher, a Swiss senior natural scientist, was really thinking applied research should perhaps be valued more than it now is. For example, one Swiss junior worried that ‘research topics with low expected/unknown society impact might not get funded’. And that ‘one tends to start thinking of projects in terms of whether they are “presentable”, as in likely to get funding…’ For example, one Swiss junior natural scientist expressed the worry that ‘It would be disastrous if competitive funding schemes would push research away from fundamental science.’ Another colleague answered to this: ‘unfortunately this is happening in selected countries’ (as indeed evidenced by e.g. Dutch natural scientists). Another natural researcher also worried about the decreasing amount of funding towards basic science. He/she thinks that we ‘should promote basic science because there is a strong, strong pressure for innovation and transaction of science, and if you kill basic science-in many countries the funding is decreasing for basic science.’ For example, there are increasing problems in Horizon 2020 funding, which ‘doesn’t recognise enough basic science in the way they rate the projects.’

And while one researcher wrote that funding might offer ‘a way for resource providers to guide research into topics that are relevant for their interests’, other Swiss natural junior researchers were quite sceptical about such developments: For example one wrote that the Swiss funder SNSF ‘definitely requires societal impact, ideally collaborations with industry, possible applications, so a lot of people in my field, including myself, have rather unrealistic writeups about super-blue-sky technology that may or may not every actually hit the ground.’

Also Dutch senior natural scientists worried about decreasing funding for basic science. Though some did see some positive aspects as well, for example that it ‘sparks creative coupling between science and industry.’ However, they also worried for example that there might be a ‘risk of over-focus on certain disciplines where societal relevance or applicability is more evident.’ Another remark was alluding to that research questions get ‘increasingly defined by applicability of the output’. One researcher wrote: ‘Over-emphasis on applied research and connections to industry. If projects/research fit, then can be a benefit. But definitely hinders very fundamental research.’ The same researcher then went on to say that it is ‘negative that nearly all 100% fundamental project funding possibilities in the Netherlands are being eliminated. Even the Science Agenda is now funded with contributions from industry.’ The national science agenda in the Netherlands had been an interesting funder experiment: An agenda for important topics for science had been established by asking the public for input. The Dutch funding agency then gave money to fund some of these agenda points. What this researcher here is worrying about is that this initiative, originally in principle disconnected from any applied aspects, now did get connected to applied aspects anyway.

Some natural science researchers succinctly emphasized the two-sidedness of collaborating with industry: ‘By having to connect with industry for funds, ability to discuss and plan research directions can be fruitful. But also very frustrating if a great fundamental question but not relevant enough for industry.’ One researcher then said that it helps to engage the industry early on in a project, not wait too long, and then there sometimes are even positive surprises what industry is interested in. Another - medical - researcher had a different but also successful strategy: to answer questions which are being posed by industry, which, this researcher said, ‘I’m not necessarily agreeing that will change the world’. But then, this researcher also makes sure to get a deal with ‘a huge chunk of money to do the things I want to do’. (med sen NL)

Simultaneously, two Dutch senior natural science researchers worried that the time horizon from research funding might in any case be too short to address the societal problems that we really would need to address: ‘Long term research is being prevented. Big societal problems require long term data.’ Another researcher fully agreed and added that ‘monitoring programs are being stopped. No incentive for scientists to continue this.’ Interesting is here that this utterance seems to relate to research that has in fact no connection to industry at all, but concerns monitoring biodiversity. So these worries are about societal problems that are not about applied science, do not necessarily need big collaborations, consortia or the involvement of stakeholders, and are not interdisciplinary projects. They are, indeed, straightforward and incremental science, just needing a long-term funding horizon. But they do carry a high societal function.

### Organization of funding: relatively free versus detailed pre-given structures

While ERC grants are relatively flexible in terms of topics and methods, consortia grants are quite pre-structured. Again, this was quite visible amongst the comments we received from researchers, with the inflexibility of consortia grants as being seen as problematic while the flexibility and autonomy provided in ERC-type grants as being appreciated. ‘[About ERC] It’s amazing, amazing the success, and this is really basic research. It’s bottom up. The researchers come with their projects. Nothing is imposed by politicians or whatever, which is not the case for the collaborative projects. So it’s really, really a fantastic institution.’ (nat sen CH)

One Swiss medical senior researcher commented that it might be positive that ‘one can guide research directions of national or international importance by specific calls.’ However, several Swiss and Dutch medical researchers emphasized that European Horizon 2020 grants might be highly problematic exactly because of this guidance. The problem with this, as researchers see it, is that this allows for researchers to impact the agenda-setting ‘before the original call comes out. Because you can influence what’s on the list.’ One Dutch researcher said that this could even be seen as a game ‘that you can play very well. And then, hopefully, it’s played by people with high integrity and not only for their own careers.’ The reason that this is possible, according to one Swiss junior medical scientist, is that ‘there are all these EU bureaucrats and they didn’t even know what to do with all the money, so they are desperate to have some professors telling them what to do with the money, and these professors then, of course, write exactly the thing that they need for their own research.’ And this is exactly what happened when this researcher eventually became part of this ‘lobbying group: There were 20 people who were phrasing this Horizon 2020 page in exactly the way so that our project would fit. This is insane.’

In addition, medical researchers do in general not value the amount of detailed proposal-writing, most of which has nothing to do with science itself: ‘these big Horizon 2020 consortia…it’s total seventy pages, such proposals, science is only four pages.’ (med sen NL). Several Swiss medical seniors also experienced that EU consortia projects in practice ‘don’t seem to work well’. These researchers compared Horizon 2020 consortia schemes to the Swiss NCCR’s and thought that analogously to the latter, European consortia are also ‘guided by particular ideas that sort out what we probably think as the most creative research.’ (med sen CH) One main underlying reason for why NCCR consortia are experienced as not working well is because they aim to not only foster high-quality science but also have other, more political, criteria attached to them. These extra criteria are then experienced as seriously complicating matters.

How this works in detail has been described by a Swiss medical senior who described the succession of several NCCR calls where extra criteria had been added only in later calls. In the beginning of these NCCR’s (first call), this researcher emphasized, ‘the intention was a very good one and it worked very well’. But then, this researcher goes on, ‘came, of course, the second and the third wave. And then people put all kinds of additional thoughts into this. Should be regional, there should be industry and there should be a very significant amount of junior funding.[…] and so it was watered down until you had so many criteria that science was just one of them. I think the system broke down.’ (med sen, CH) When asked, this researcher emphasized that it was not the amount of money itself that was problematic here (this first call ‘transformed research, no doubt’), but that the problem was that due to political pressures other aspects than science started playing a role as well: ‘the money was significant enough that politicians became interested in this. And from that point on, I would simply say it was watered down.’ The problem, according to Swiss senior medical scientists, is that there was too little academic freedom left, there was in the end ‘Too much other influences outside of the science.’ Also a natural scientist worried that the funding situation in Switzerland might get worse due to such pressures to justify funds: ‘Unfortunately, because of all the pressure around us, we’re also going downhill…. There is a lot of pressure to be more competitive, to add more around it, to justify the funds. I think the politicians don’t always understand how the science works.’ (nat sen CH)

The problem, according to Swiss senior medical researchers, occurs if you add too many other aspects aside from the science itself, aside from aspects regarding scientific excellence. If you try to serve too many agenda’s, ‘then you are nowhere. Then you don’t know where your attention is.’ And then it goes wrong, according to those researchers, even though they do understand and value the reasoning behind it: that due to the big amount of money put into such NCCR’s ‘obviously, people look at this very carefully.’ And that then politicians think ‘if this is so much money, you have to fulfil, at least, five secondary roles as well.’ But then, these researchers perceive, matters go into the wrong direction, at least in terms of science. A Swiss junior natural scientist emphasized that in general one would need to ‘reduce the number of boxes you have to tick, because if you want to do everything then you achieve nothing.’ And another Swiss natural scientist said that EU projects are ‘almost not worth the money you get. I have too many of those.’ In his/her eyes, the main problem is ‘The amount of work you have around with managing it…’

Another issue are smaller interdisciplinary Swiss Synergia projects, according to Swiss senior medical researchers, which are small consortia of three to four members. These work well in the eyes of the medical senior interviewees. ‘The Synergia is very focused. I think this has never run into the secondary problems [that NCCR’s and EU consortia have, according to these medical researchers].’

Interestingly, one senior Swiss natural scientist with personal experience of both NCCR’s and ERC’s was very happy with NCCR’s, because they can ‘give you [the individual researcher] time’ to venture into new fields, try out new things. The conditions under which this can happen, this researcher emphasized, is if the director lets you do so, provides the researchers with sufficient autonomy, if he/she says: ‘You just use your money however you want to use it.’ Also Dusdal and Powell (2021) emphasized that the person in charge of the network has an important function to shape the collaboration.

## Discussion

We find that researchers prefer ERC-type funding not per se due to the innovation or excellence component, but because several aspects of the funding specifics align mostly faithfully with how they experience science should effectively be done, also in terms of impact.

First, researchers across groups experience that science conducted in big consortia networks does not seem to work well in practice: the networks are too loose to be effective, and members might push into such structures who do not help but instead might even decrease the quality of the resulting science. In addition, individual members might not do their best in such bigger teams, perhaps due to ownership issues. There is also clearly added bureaucracy and communication problems across different groups in such large multi-disciplinary and international groups (see also Dusdal and Powell 2021 for similar findings). Even in smaller teams, the epistemic distance between members from different disciplines can provide substantial challenges (Stephens and Stephens 2021). Researchers in our study have not reported the same types of troubles occurring in ERC-like types of teams that work closely together and are epistemically more aligned. It is also likely that the PI in an ERC grant acts as an anchor point around which all actions are concentrated. What is more, the PI also has the flexibility to choose team members that would function socially within the team. Several recent papers have outlined how important such social aspects are to do good collaborative work (Dusdal and Powell 2021; Hesjedal 2022).

Secondly, and perhaps most importantly, researchers across groups are highly skeptical of impact-driven funding schemes, such as EU consortia funding (but also other national ones). Medical researchers experience that such short-term impacts are essentially highly unrealistic in terms of their time horizon. Natural researchers express that some of the most socially valuable scientific work would simply need a long-time horizon and not a big network to perform (such as monitoring). Medical researchers point out that involving stakeholders is in practice experienced as very difficult, both because it is not always clear how this should happen and what they could contribute, but also because it can have adverse effects of too little time to do the core work. This can result in overhasty implementation. Indeed, also other studies have highlighted such challenges, suggesting that one would need to analyze in which cases it does indeed pay off (Lemos et al. 2018; Wyborn et al. 2019). Some of our researchers told us how they flexibly deal with such challenges in practice: For example, natural researchers experience that industry can have very different goals, but that it can work out if the collaboration is done with a lot of care and communication. One medical researcher said that it can work out to simply perform what industry wants and then keep some of the money to do interesting fundamental work on the side. Finally, medical researchers highlight that real impact cannot be planned in such manner – real scientific breakthroughs are essentially unpredictable, and findings can pay off only decades later (see also Copeland 2019). What this all often results in is that researchers have the feeling to have to lie, to have to overpromise, regarding impact in grant proposals. They are ‘hot air’. One researcher suggested that the only thing that could be done to foster impact would be to reward researchers with a certain personality of being curious, working diligently, and playing around. In essence, that sounds more like the rewards provided via a curiosity-driven and personal ERC grant. In addition, many natural science researchers were highly worried about basic funding receiving too little of its share, both in national and in EU funding; current funding schemes even being a threat to fields that are more fundamental.

Third, researchers experienced that the pre-structuring of consortia-type funding schemes is not valuable. For example, while medical researchers appreciate that specialized calls might enable policy makers to target science towards solving specific problems, the process in which such calls in EU funding are being made in practice is experienced as too biased. Several of our medical interviewees, who had insider experience, said this is essentially like ‘a game’, in which researchers may even tailor calls to suit their own needs. Also, the types of detailed proposals for EU consortia calls are clearly not being appreciated; a large part of these proposals not even having anything to do with the proposed science. And even though several of our interviewees can understand why politicians would feel forced to make big-budget science more accountable by adding further elements, both medical and natural science researchers experienced that such added elements (beyond scientific ones) were overall distractive and complicated matters. One researcher said that apparently politicians don’t always understand how science works. Another researcher put it as such: ‘If you want to do everything you achieve nothing.’ Consequently, this researcher (and others) suggested that the less boxes you must tick in a funding application the better for the resulting science. Funding that allows for more autonomy and flexibility was clearly experienced as more valuable by most of our interviewees – and this could even happen in bigger structures if the person in charge allows for it.

Against the arguments of other science studies colleagues (Scholten et al. 2021; Falkenberg et al. 2022), we argue that it might be most valuable to channel even more money towards ERC-types of funding, thus reducing the adverse effects of competition, rather than pulling budget over to other types of more top-down funding schemes. Our suggestion is in line with findings showing that research may thrive better if researchers are provided with sufficient autonomy, including the possibility to play around (Laudel 2006). Indeed, scientists may value the ERC precisely because, as Roumbanis (2019) puts it, many universities in Europe ‘have taken on a more market-oriented approach that has changed the core of academic work life.’ In this context, the need for ‘protected spaces’ (Laudel 2017) in which scientists can work on meaningful research for which they are intrinsically motivated seems higher than ever. Also the KNAW (2019) argues that the Dutch research funder NWO should (re)tailor more of its funding for such types of free science and away from agenda-driven science (and importantly, there was very recently a political decision to fund more investigator-driven science in the Netherlands). Even with regards to the UK REF, experts start appreciating that “pre-conditions for such [research] governance include intellectual freedom in research” (Oancea 2019).

Importantly, also other recent evidence shows that a freer investigator-led approach does not preclude addressing applied or problem-generated topics, such as climate change or clean oceans. For example, a 2018 evaluation report of ERC projects showed that nearly half of the funded projects already have a societal impact, while around 75% are predicted to do so in the longer term - and that without societal impact being a criterion of selection (European Research Council 2019; this evaluation was assisted by independent experts selected by the ERC). The report also showed that many ERC projects are strongly interdisciplinary, with around 70% of the evaluated projects having led to results applicable to other areas of research, while around 60% of them brought together two previously rather unconnected research areas.

Scientists in our study also experienced very positive effects from Swiss Sinergia funding, which provides relatively free types of funding for interdisciplinary small-scale projects (see also Ayoubi et al. 2019). Our findings suggest that such types of teams may cultivate an optimal form of focused collaboration, which Hanson (2018) called ‘disciplined collaboration’ in the business world. As such, scientific collaboration might work best if it steers a middle course between under-collaboration (isolation) and over-collaboration (unnecessarily complex forms of cooperation that have a negative effect on work performance), with disciplined collaboration producing the best and most effective results. Arguably, also ERC-type funding leads to such disciplined collaboration.

That researchers in general are already overworked, lacking time and feeling that they have little ‘space to maneuver’ (Åm 2019; see also Sigl et al. 2020) is an often-heard complaint increasingly being made by researchers (see e.g. Wellcome Trust report 2020). Interestingly, Åm (2019) cites older scientists who feel that there was simply more space and freedom for good discussions in earlier decades, and that this by itself led to an increased degree of reflexivity. Åm further advises that effectively incorporating aspects of responsible research and innovation only works in practice if such space and freedom (again) is provided, and that it in addition is mandatory that scientists develop a sense of ownership of such concepts. These ideas integrate well with our own findings on the need for focus, time, freedom and ownership – and that this could ultimately lead to better research also in a societal sense.

In conclusion, we suggest that it is important to rethink the recent international drive towards multidimensional consortia funding schemes. Our study suggests that it might be more important to invest more in investigator-led ERC-types of science, and that this might not be to the detriment of societal involvement and relevance (see also KNAW 2019, 2020). It seems that what researchers nowadays increasingly lack is the time and the necessary academic freedom to focus their work in the most efficient ways.

## Acknowledgements

First of all, we would like to thank our interviewees in the Netherlands and Switzerland for taking the time to share their insights with us. We would also like to thank the Swiss Academies of Arts and Sciences, and in particular Roger Pfister, for hosting the sessions in Switzerland. We are also indebted to Gerd Folkers, who guided us and provided contacts to Swiss scientists. We also thank the Royal Netherlands Academy of Arts and Sciences, and Jean Philippe de Jong in particular, for helping to contact interviewees, hosting the discussion sessions in The Netherlands and engaging the Swiss Academies in the project. We also thank Jean Philippe for his comments and edits on a previous version of this manuscript. Furthermore, we would like to thank Jeannette Pols, Barend van der Meulen and Peter van Hoesel for good advice throughout our project, as well as Danny van den Boom with helping to make the Meetingsphere sessions a success.

## Footnotes

1 In the Results, quotes are indicated. Where not obvious from the surrounding text, these quotes are accompanied by the following abbreviations to signalize who made the quote: med = researcher in a medical field; nat = researcher in a natural science field; jun = junior; sen = senior; NL = researcher currently based in the Netherlands; CH = researcher currently based in Switzerland

